# Subtype-specific downregulation of voltage-gated sodium channels shapes neuronal responses to neuroinflammation

**DOI:** 10.64898/2026.08.24.746613

**Authors:** Daniela Jacobsohn, David Guenoun, Nathalie Hertrich, Pascal Fenske, Sabrina Pommer, Shyamala Mani, Angela M. Kaindl

**Author notes:** These authors contributed equally (Daniela Jacobsohn, David Guenoun, Shyamala Mani, Angela M. Kaindl). Corresponding author (A.M. Kaindl).

## Abstract

Epilepsy is one of the most common neurological disorders, affecting more than 50 million people worldwide. Among the genetic etiologies of epilepsy, variants in genes coding for ion channels are vastly represented and characterized. Notably, loss-of-function (LoF) mutations in voltage-gated sodium channels (Na_V_) genes can result in a wide range of phenotypes including West syndrome, autism spectrum disorder, or Dravet Syndrome. Although the implication of Na_V_ subtypes in epileptic syndromes and the relationship between seizures and inflammation have been extensively described, subtype-specific neuronal responses to inflammation in the context of Na_V_ loss-of-function remain poorly understood. In this study, we investigated the consequences of subtype-specific downregulation of Na_V_ expression in primary mouse cortical neurons. Using shRNA-mediated silencing of *Scn1a*, *Scn2a*, or *Scn8a*, we generated neuronal cultures with reduced expression of Na_V_1.1, Na_V_1.2, or Na_V_1.6 and evaluated neuronal survival, inflammatory gene expression, and global transcriptomic responses under basal conditions and following an inflammatory challenge. Subtype-specific Na_V_ downregulations did not produce a uniform phenotype. Rather, minor differences under basal conditions led to important discrepancies following exposure to an inflammatory stimulus. Notably, Na_V_1.1 reduction was associated with synaptic transcriptional changes, whereas Na_V_1.6 downregulation led to a substantial inflammatory signaling remodeling. Our observations suggest that the consequences of Na_V_ dysfunction are not only determined by their role in neuronal excitability but also depend on subtype-specific responses to inflammatory cues. They notably shed light on the relevance of inflammatory events in the onset and progression of epileptic syndromes related to Na_V_ loss-of-function mutations.

## Introduction

Epilepsy is one of the most common neurological disorders, affecting more than 50 million people worldwide (1). It encompasses a broad spectrum of etiologies, including genetic, structural, auto-immune, infectious or of undetermined origin. Genetic factors play a particularly important role in many epilepsy syndromes, ranging from complex polygenic susceptibility to monogenic disorders caused by pathogenic variants in individual genes. Since the first identification of *CHRNA4* as the first gene associated with a monogenic epilepsy syndrome in 1995, more than 1500 genes have been associated with epilepsy (2). Among these, genes coding for ion channels represent one of the largest and best-characterized groups, responsible for disorders collectively referred to as channelopathies. This is notably the case of mutations occurring in voltage-gated sodium channels (Na_V_s) in which loss (LoF) or gain of function (GoF) variants can cause epilepsy and neurodevelopmental disorders. In particular, LoF variants affecting Na_V_s can result in a wide range of phenotypes including West syndrome, autism spectrum disorder, or the prototypical developmental and epileptic encephalopathy (DEE), Dravet Syndrome (DS) (3).

Na_V_s are a group of nine sodium channels expressed at the membrane of neurons in both the central nervous system (CNS) and the periphery. They are composed of a main α-subunit that forms the ion conducting pore and voltage sensor of the channel, associated with smaller β-subunits that modulate channel activation and gating. In response to membrane depolarization, Na_V_s generate an inward sodium current crucial to the generation and propagation of action potentials. In the mammalian CNS, four Na_V_s, Na_V_1.1, Na_V_1.2, Na_V_1.3, and Na_V_1.6, are predominantly expressed in neurons, with distinct developmental and neuronal-type specific patterns. Na_V_1.3, encoded by *SCN3A*, is highly expressed prenatally but declines substantially after birth. In contrast, the expression levels of Na_V_1.1 (*SCN1A*) and Na_V_1.6 (*SCN8A*) increase after birth, whereas Na_V_1.2 (*SCN2A*) remains stable across developmental stages (3). These differences are particularly functionally relevant in regard to their specific neuronal distributions: Na_V_1.2 and Na_V_1.6 are mostly expressed in excitatory neurons, and Na_V_1.1 is, in adults, enriched in inhibitory interneurons, notably in the hippocampus. Consequently, loss of Na_V_1.1 preferentially compromises the excitability and function of inhibitory interneurons, whereas reduced NaV1.2 or NaV1.6 function primarily affects excitatory neuronal activity, providing an important basis for the distinct clinical consequences associated with pathogenic variants in *SCN1A*, *SCN2A*, and *SCN8A*.

Similarly, Na_V_s exhibit distinct subcellular distributions within neurons, notably at the axon initial segment (AIS), a specialized structure of the proximal axon that plays a critical role in the initiation of action potentials. Na_V_1.1 is predominantly present in the soma and in the proximal region of the AIS, and Na_V_1.2 in the proximal part of AIS, the soma and in dendritic compartments. While during early neurodevelopment, Na_V_ 1.6 is mostly concentrated in the distal region of the AIS, it becomes, as maturation and myelination occur, the dominant Na_V_ all along the AIS and at the nodes of Ranvier (4). These spatially and temporally distinct expression patterns contribute to the specialized roles of individual Na_V_s in regulating neuronal excitability, action potential initiation, and propagation.

Beyond their canonical role in electrical signaling, increasing evidence suggests that neuronal ion channels may also influence cellular responses to injury and immune-mediated stress. Neuroinflammation occupies a central position in the pathophysiology of the onset and/or chronicity of neurological and neurodevelopmental diseases, including epilepsy where the close relationship between seizures and neuroinflammation is well established. Indeed, while neuroinflammation is mainly driven by microglial reactivity, neuronal dysfunction and damage can lead to the onset of neuroinflammatory processes (5–7). Furthermore, neurons have been shown to be able to act as immunomodulators and release cytokines (8–10). In return, inflammation tends to increase neuronal excitability through the release of inflammatory cytokines such as interleukin (IL)-1β, tumor necrosis factor (TNF)-α, or IL-4 (11). Therefore, neuroinflammatory mechanisms, through this concept of a self-reinforcing vicious cycle, have long been hypothesized as a contributor to both disease progression, high-seizure frequency and drug-resistance in DEEs (12,13). These observations are at the core of the rationale behind the use of steroids in drug-resistant forms of epilepsy, and support the use of cannabidiol (FDA approved in 2018) in the treatment of DS (14).

Importantly, literature suggests that Na_V_ dysfunction can modulate neuroinflammatory responses. Mouse models of DS have demonstrated increases in microglial activation and inflammatory markers, and, conversely, maternal immune activation has been reported to reduce *Scn1a* expression, providing evidence that inflammatory processes can directly influence Na_V_ expression (15–17). In addition, Qiao *et al*. demonstrated that the decrease in Na_V_ expression following status epilepticus is correlated with the subsequent neuronal loss (18). Together, these observations raise the possibility that Na_V_ dysfunction may not only alter neuronal excitability but also modify the capacity of neurons to respond to neuroinflammation.

However, whether reduced expression of individual Na_V_s differentially affects neuronal responses to inflammation remains poorly understood. Addressing this question is of particular relevance to genetic epilepsies, in which pathogenic variants affecting different Na_V_s can produce markedly distinct phenotypes despite their shared involvement in the regulation of neuronal excitability. Understanding how the selective LoF of Na_V_s differentially affects the neuronal response to inflammation as well as which pathways these processes imply may therefore provide new insights into the cellular mechanisms underlying these disorders and identify potential therapeutic targets.

In this study, we investigated the consequences of specific reduction in Na_V_ subtypes expression in primary mouse cortical neurons. Using viral vectors encoding short hairpin RNAs (shRNAs) targeting *Scn1a*, *Scn2a*, or *Scn8a*, we selectively reduced the expression of Na_V_1.1, Na_V_1.2, or Na_V_1.6 and assessed neuronal survival, inflammatory cytokines secretion, and transcriptomic responses under basal conditions as well as following an inflammatory challenge. We hypothesized that reduction of distinct Na_V_ subtypes would lead to different neuronal responses to inflammation, thereby revealing specific mechanisms linking Na_V_ dysfunction to neuronal vulnerability and inflammatory signaling.

## Materials and Methods

### Primary neuronal cultures

Primary cortical neuron cultures were established using *C57BL/6 J* mice at age P0-P1 and following the protocol established by Beaudoin *et al.* (19). The cortices were first extracted from mice and the hippocampus and meninges removed. Samples were then incubated with 0.25% trypsin EDTA (Santa Cruz Biotechnology, Inc., sc-391060) for 12 minutes before addition of FBS and treatment with 0.5mg/ml DNAse 1 for 4 min. Afterwards, the samples were mechanically dissociated by successive pipetting through a blunted glass pipette. Samples were then resuspended in DMEM with 10% FBS and incubated for 2 hours before changing the medium to Neurobasal™-A (Gibco, 10888022). Neurons were cultured on 14mm glass coverslips coated with 0.02 mg/mL poly-DL-ornithine hydrobromide (Sigma, P8638) in 24 Well cell culture dishes at a density of 500,000 cells/ml. At DIV 2 neurons were treated with 0.5µM of cytosine arabinoside for 24h. After 24 hours of incubation, the medium was changed to Neurobasal™-A. All experiments were performed at days *in vitro* (DIV) 10.

### Microglial cultures

Microglia cultures were established using P4 C57BL/6J mice according to the protocol described by Platow *et. al* (20). Whole cortices were extracted and meninges removed. Brain tissue was dissociated into small fragments and incubated with Trypsin 2.5% (Gibco,15090046) for 25 min. 0.1 mg/ml of Deoxyribonuclease I (DNase 1) (Sigma, DN25) was then added and samples were centrifuged at 1,200g at 4°C for 5 min. They were resuspended in Dulbecco’s Modified Eagle Serum (DMEM, Gibco) with 10% fetal bovine serum (FBS) and 1% penicillin-streptomycin (P/S) (DMEM-C) and seeded in culture flasks for 12 days. At DIV 12 microglia were plated in 6 well plates at a density of 300,000 cells/ml in DMEM-C. On DIV 14 cells were stimulated with lipopolysaccharide (LPS, E. coli serotype O111:B4) 10 µg/mL for 24h after which the medium was changed to neuronal culture medium. Neuronal culturing Medium Neurobasal™-A Medium with 1% B-27™ Supplement (50x), 0.5% P/S and 0.5 % GlutaMAX™ Supplement (Gibco, 35050061) was used. After 24h the neuronal culture medium was taken off and filtered using a Millex-GV Filter, 0.22 µm (Millipore) to be later used as an inflammatory environment (CM-inf) for neuronal cultures. CM-inf was then stored at -70°C until use. For control conditions (CM-Ctl), microglia were cultured following the same protocol but were not stimulated with LPS. Successful stimulation was confirmed by rt-qPCR targeting prostaglandin-endoperoxide-synthase (PTGS2) and tumor necrosis factor-α (TNF-α) in microglia. Similarly, pro-inflammatory cytokine enrichment of CM-inf was confirmed using cytokine arrays (**Supplementary Fig. 1**).

### Subtype-specific downregulation of Na_V_s

To downregulate the expression of specific Na_V_ subtypes in our neuronal cultures, cells were transduced with a lentiviral construct carrying short hairpin RNAs (shRNA) sequences complementary to either *Scn1a*, *Scn2a* or *Scn8a* and mCherry fluorescent reporter sequence to allow for the confirmation of successful transduction. A scrambled (Scr) sequence was used as control for all experiments. ShRNA sequences are detailed in **Table 1**.

**Table 1:**
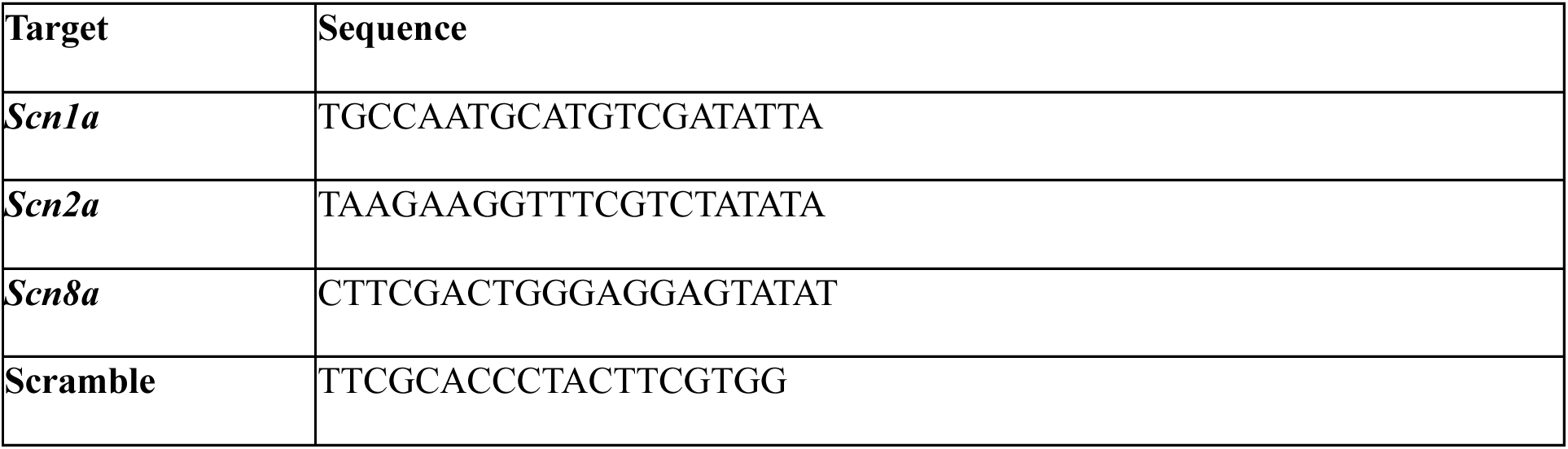
ShRNA sequences.

### Stimulation

At DIV 9, Neurons were stimulated using a 1:10 dilution of CM-Inf in NBA-C as described above. 24h after stimulation, neurons were either directly frozen and stored at -70°C for RNA extraction or fixed for immunocytochemistry using 4% Paraformaldehyde with 4% Sucrose Solution for 15 minutes.

### *In vivo* channel rt-qPCR

For each sample, a *C57BL/6 J* mouse at P0, P7, P14 or P56 was sacrificed. RNA was extracted using NucleoSpin RNA Plus XS (Macherey-Nagel, 740990) and channel expression was quantified by rt-qPCR as described below.

### Immunocytochemistry (ICC)

Staining was done using the protocol described by Platow *et al.* (20). Coverslips were briefly washed with PBS+ 0.5% bovine serum albumin (BSA) + 0.5% Triton X before permeabilization with Triton X 0.05% for 15 minutes and blocking using 10% Donkey Serum (DS) + 5% BSA for 30 minutes. The samples were then incubated with the primary antibody (**Table 2**) diluted in the blocking solution overnight. After washing with PBS, they were incubated with the secondary antibody for 2 h. They were then washed again with PBS and incubated with DAPI (BD Pharmingen™, 564907, 1:1000), for 20 min. Coverslips were then washed and mounted on Glass Slides using Immu-Mount™ (Epredia™, 9990402). Images were captured using a Zeiss Spinning disc confocal microscope (CSU-X1, Yokogava) controlled by the ZEN blue 2012 software (2.6) with a 20x air objective (Zeiss Objective Plan-Apochromat 20x/0.8 M27) and a Zeiss EMCCD camera (QImaging Rolera em-c2, 1004×1002 pixels). Image analysis was performed using Fiji (ImageJ) version 2.14.0 Software (21). For each replicate at least ten fields of view were imaged, with > 500 cells analyzed for each replicate.

**Table 2:** List of primary antibodies used in ICC.

|  | Dilution | Manufacturer | Reference |
| --- | --- | --- | --- |
| Anti-Map2 | 1:2500 | Abcam | #ab5392 |
| Anti-Gad67 | 1:500 | MAB5406 | Sigma Aldrich |
| Anti-Nav 1.1 | 1:500 | Sigma Aldrich | AB5204 |
| Anti-Nav 1.2 | 1:500 | NeuroMab | 75-024 |
| Anti-Nav 1.6 | 1:5000 | Sigma Aldrich | WH0006334M4 |

### Reverse Transcription quantitative PCR (RT-qPCR)

RNA was extracted using the NucleoSpin RNA XS Kit (Macherey-Nagel, 740990) following the protocol issued by the manufacturer. Extracted RNA was then reverse transcribed into cDNA using the iScript™ cDNA Synthesis Kit (Bio-Rad, 1708890). For qPCR, primers were used together with iTaq Universal SYBR Green Supermix (Bio-Rad, 1725120). Primer sequences are displayed in **Table 3**. After stimulation levels of *Cxcl10*, Il-*6* and *Ifn*-β were determined. RT-qPCR results were normalized to the housekeeping gene Rpl13a. Analyses were performed using the StepOne software V2.3 (Thermo Fisher Scientific, Waltham, MA, USA) and a relative quantification approach was used, according to the 2-ddCT method. For the histograms the average 2-ddCT value for each well was plotted.

**Table 3:**
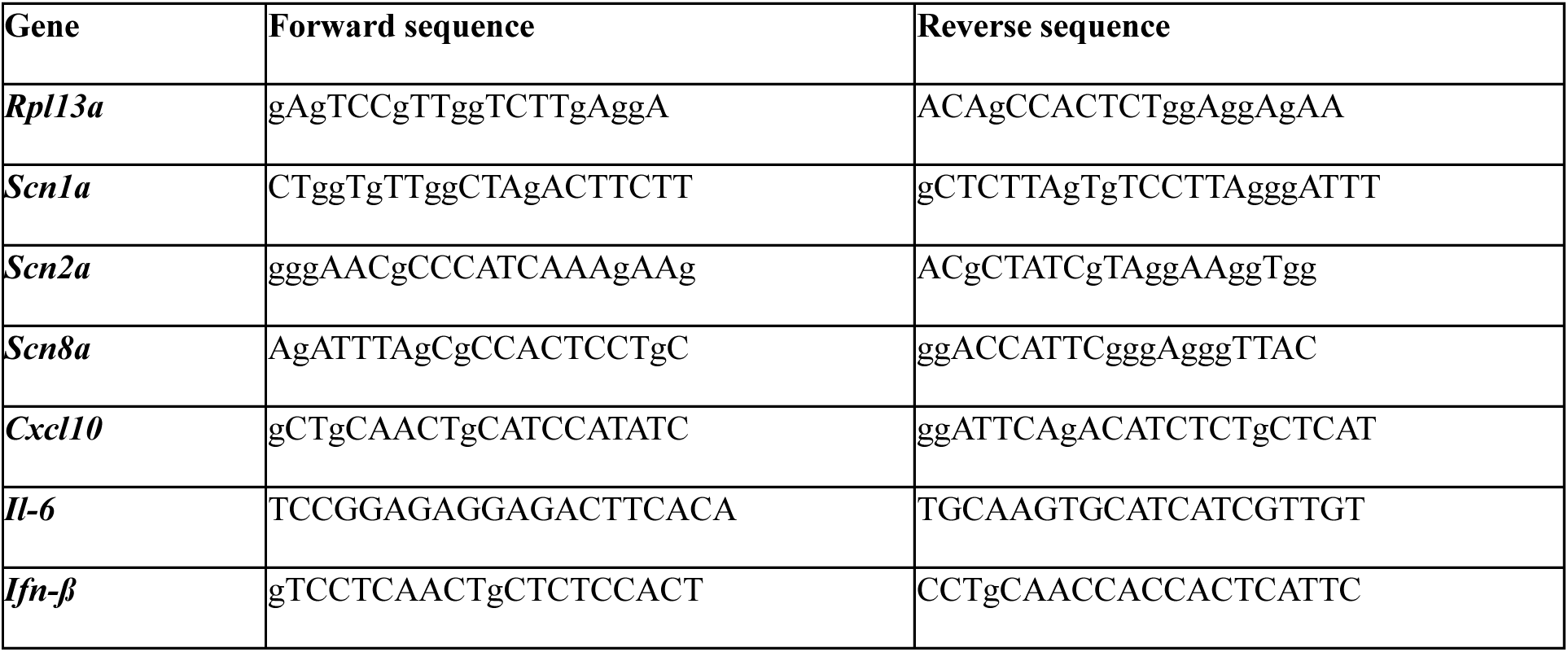
Details of primers used in RT-qPCR.

### Western Blot

Proteins were extracted using the Mem-PER™ Plus Membrane Protein-Extraction-Kit (89842, Thermo Fisher Scientific) following the manufacturer’s instructions with following modifications: Step 1.3. was skipped, 200 – 300 µL permeabilization buffer and 25 -50 µL solubilization buffer were used for each condition/well, protease and phosphatase inhibitor cocktail 1:100 (Sigma, PPC1010-1ML) was added to the buffers. Samples were denatured at 90 °C during 5 min before migration in an 8 % percent tris-acetate gel. After migration, proteins were transferred onto a nitrocellulose membrane (88018, Thermo Scientific™). The membranes were washed with PBS-T, blocked with 5% BSA (Sigma-Aldrich, A9647) in PBS at room temperature and incubated overnight at 4 ℃ with the primary antibody (**Table 4**). Subsequently, the membranes were washed at room temperature and incubated in goat anti-mouse immunoglobulins/HRP (Dako, P0447) and donkey anti-rabbit IgG (H+L) at room temperature for 1.5 hours. Both secondary antibodies were diluted 1:10 000 in PBS. After washing, ECL (Bio-Rad, 170-5060) was added before acquisition using an Azure 600 Imaging System (Biozym). Analyses of the acquired images were performed using Fiji (ImageJ) version 2.14.0 Software (21).

**Table 4:** List of primary antibodies used in Western Blot.

|  | Dilution | Manufacturer | Reference |
| --- | --- | --- | --- |
| **Anti-Calnexin** | 1/5,000 | Sigma-Aldrich | #C4731-.2ML |
| **Anti-Na<sub>v</sub> 1.1** | 1/5 | DSHB | K47/715 |
| **Anti-Na<sub>v</sub> 1.2** | 1/200 | NeuroMab | 75-024 |
| **Anti-Na<sub>v</sub> 1.6** | 1/500 | Alomone | ASC-009 |

### Transcriptomics

Three biological replicates were used for transcriptomics. Purified RNA, extracted as specified above, was thawed and processed according to the instructions of the AccuraCode® RNA-Seq kit (Singleron Biotechnologies GmbH, 10710174) with the following modifications. Briefly, polyadenylated mRNA from different samples was captured using paramagnetic AccuraCode bulk beads (Singleron Biotechnologies GmbH, 1200030080). During reverse transcription, the mRNA was converted into cDNA and labeled with sample-specific barcodes and unique molecular identifiers (UMI). The labeled cDNA was pooled, amplified and used to construct RNA-seq libraries according to manufacturer’s instructions (AccuraCode® RNA-Seq kit, Singleron Biotechnologies GmbH, 10710174). The libraries were sequenced on an Illumina NovaSeq X instrument using a paired-end 150-bp approach by Macrogen Europe (Amsterdam).

### RNA-seq data analysis

Raw reads were demultiplexed according to the well barcodes using CeleScope v2.7.4 multi_bulk_rna (www.github.com/singleron-RD/CeleScope; Singleron Biotechnologies GmbH) and the individual R2 fastq files run through the nf-core/rnaseq pipeline (v3.14.0) using the STAR Salmon for mapping and quantification to generate the gene count tables.

### Post processing of Transcriptomics data

Normalization of counts matrixes and differential expression analyses were done using the CeleLens Cloud software (Singleron). mRNAs with adjusted pval <0.05 and absolute fold-change ≥ 2 were considered differentially expressed (DE). Morpheus (https://software.broadinstitute.org/morpheus) was used to generate heatmaps and for hierarchical clustering using the one minus Pearson coefficient method. Pathway enrichment analyses within clusters were done using WebGestalt (22). Gene Ontology (GO) (http://geneontology.org, 2019.05) and Kyoto Encyclopedia of Genes and Genomes (KEGG) (http://www.kegg.jp/, 2019.05) pathway enrichment analyses were performed respectively. Data visualization was performed in R version 4.4.1 (23) using the following packages: ggplot2 (24) and pheatmap (25).

## Statistics

Data are presented as mean ± standard deviation (SD). Normality of distributions was systematically assessed using the D’Agostino & Pearson test. For comparisons between two groups, we used either Student’s t-test (for normally distributed data with equal variances) or the Mann–Whitney U test (for non-normally distributed data). For multiple group comparisons, two-way ANOVA was applied when assumptions of normality and homoscedasticity were met, followed by Fisher’s least significant difference (LSD) post hoc test. In cases where assumptions were not fulfilled, non-parametric alternatives such as the Kruskal–Wallis test were employed, followed by Dunn’s post hoc test. Outliers were removed using ROUT method with Q = 1%. Exact p-values are reported throughout, and a threshold of p < 0.05 was considered statistically significant.

## Results

### Expression and localization profiles of Na_V_ subtypes in primary cortical neurons follow key features observed *in vivo*

To better characterize our primary mouse cortical neuronal cultures, we first assessed the expression patterns of Na_V_s using rt-qPCR at three different time-points: a) when establishing the cultures at 1 day *in-vitro* (DIV), b) at DIV7, and c) at DIV10, our chosen time-point for further experiments. In all time points, *Scn2a* was the most expressed Na_V_ gene, and both *Scn1a*/*Scn2a* and *scn8a*/*Scn2a* ratios decreased over time (**Fig. 1a, b, c**). We then examined how these patterns of expression related to *in vivo* expression of Na_V_s by analyzing cortex lysates of mice at post-natal day (P)0, P7, P14 and P56. When compared to their P0 expression, all three genes showed increased expression levels over time with regular increases from P0 to P56. Similarly to what was observed *in vitro*, *Scn2a* displayed the highest expression levels, followed by *Scn8a* and *Scn1a* at all time points (**Fig. 1d**). However, while *in vitro Scn8a*/*Scn2a* and *Scn1a*/*Scn2a* ratios diminished over time, they respectively remained stable and increased over time *in vivo* , highlighting the already reported post-natal expression increase of *Scn1a* expression.

**Figure 1:**
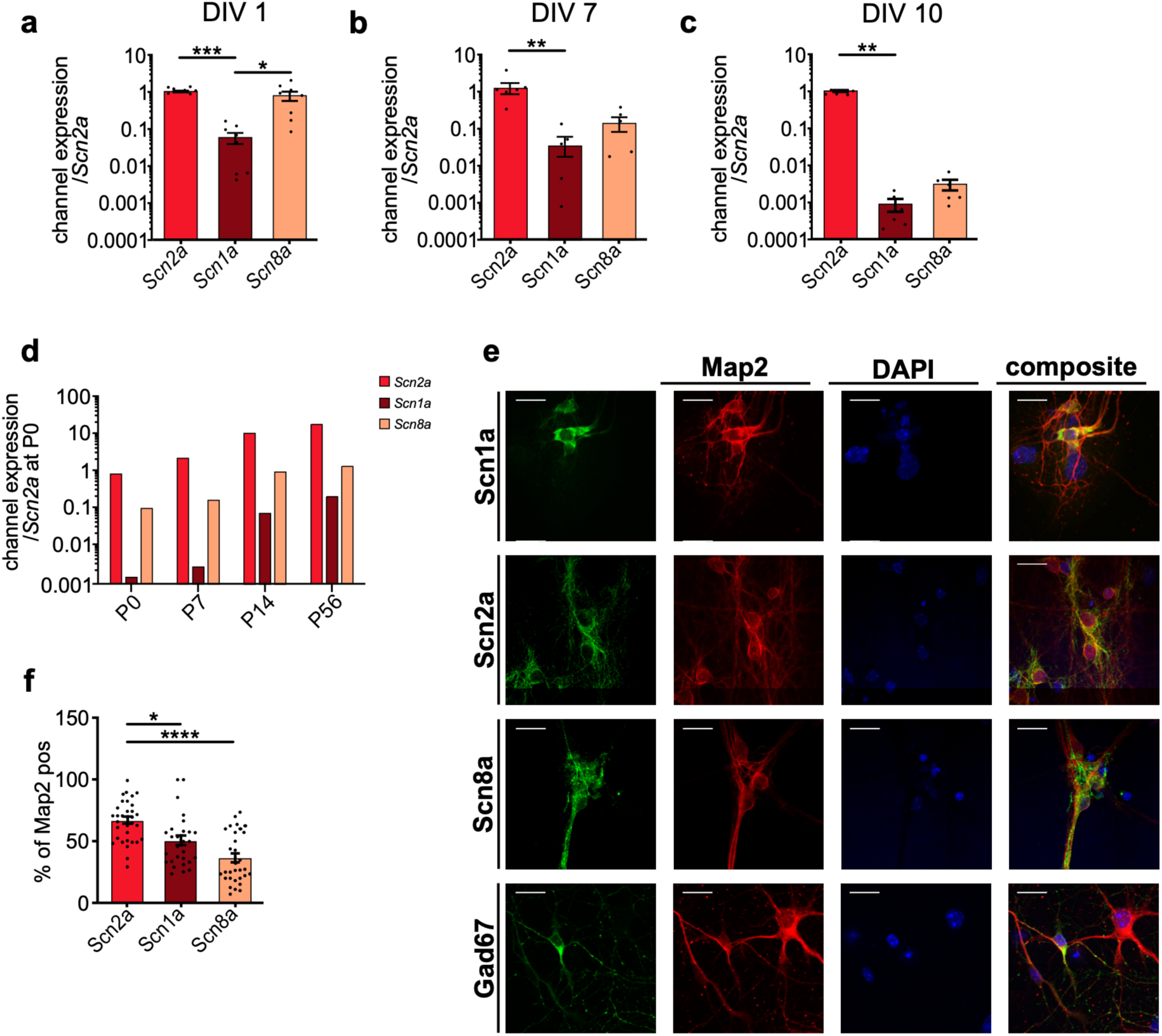
Expression and localization profiles of Na_V_ subtypes in primary cortical neurons follow key features observed in vivo. RT-qPCR results of the analyses of the expression of Na_V_ subtypes *in vitro*: **a.** at DIV 1, **b.** at DIV 7, **c.** DIV10, d. *in vivo* expression of Na_V_ subtypes at P0, P7, P14 and P56 normalized to the expression of *scn2a* at P0. **e.** Representative immunocytochemistry of primary neuronal cultures at DIV10 marked with Map2 (red), DAPI (blue), and either Na_V_ 1.1, Na_V_ 1.2, Na_V_ 1.6 or Gad67(green) **f.** Proportion of Map2^+^ cells colocalized with Na_V_ 1.1, Na_V_ 1.2 or Na_V_ 1.6.

Second, we used immunocytochemistry to describe the neuronal populations present in our cortical cultures as well as the localizations of Na_V_s. Using the neuronal marker Map2, we first assessed the proportion of neurons expressing the Na_V_s. In our analysis, 66.89% of neurons expressed *Scn2a*, 49.90% expressed *Scn1a* and 35.61% expressed *Scn8a* (**Fig. 1e, f**). An average of 36.0% of neurons were positive for the inhibitory neuron marker Gad67 among all MAP2^+^ cells across our cultures, highlighting the presence of both excitatory and inhibitory neuronal populations and coherent with the literature (26). Although *Scn1a* expression was enriched in GAD67^+^ neurons, it was not exclusively restricted to inhibitory neurons, in agreement with previous *in vivo* observations (27,28). As expected, *Scn2a* and *Scn8a* expression was detected across both Map2^+^/Gad67^+^ and Gad67^-^ neuronal populations.

Finally, we examined the subcellular localizations of Na_V_1.1, Na_V_1.2 and Na_V_1.6. Consistent with their previously described localizations *in vivo*, Na_V_1.1 was predominantly localized in the soma and the proximal regions of neurites, whereas Na_V_1.2 and Na_V_1.6 were detected in the soma and all neuronal processes (**Fig. 1e**).

Together, these results indicate that our primary mouse cortical neuronal cultures reproduce key features of the spatial and temporal expression patterns of Na_V_ subtypes *in vivo* and therefore provide a suitable model to investigate the consequences of subtype specific Na_V_ downregulations.

### shRNA-mediated downregulation selectively reduces Na_V_ subtypes expression

We next sought to establish an experimental model of subtype-specific Na_V_ downregulation. Primary cortical neurons were transduced with lentiviral vectors encoding complementary short hairpin RNAs (shRNA) targeting *Scn1a* (ΔScn1a)*, Scn2a* (ΔScn2a), or *Scn8a* (ΔScn8a) with a scrambled shRNA sequence (Scr) used as control. RT-qPCR results confirmed a significant reduction in the expression of all targeted Na_V_ subtypes in each condition compared to Scr. *Scn1a* expression was reduced by 61.5% (p = 0.015) in ΔScn1a, while *Scn2a* and *Scn8a* expression were reduced by 65.1% (p = 0.0002) and 50.8% (p = 0.0043), in ΔScn2a and ΔScn8a respectively (**Fig. 2a-c**). These reductions were further confirmed at the protein level by western blotting with Na_V_1.1, Na_V_1.2 and Na_V_1.6 protein levels respectively reduced by 43.2%, 25.6% and 48.0% in each condition (ΔScn1a, ΔScn2a or ΔScn8a) when compared to Scr (**Fig. 2d-i**).

**Figure 2:**
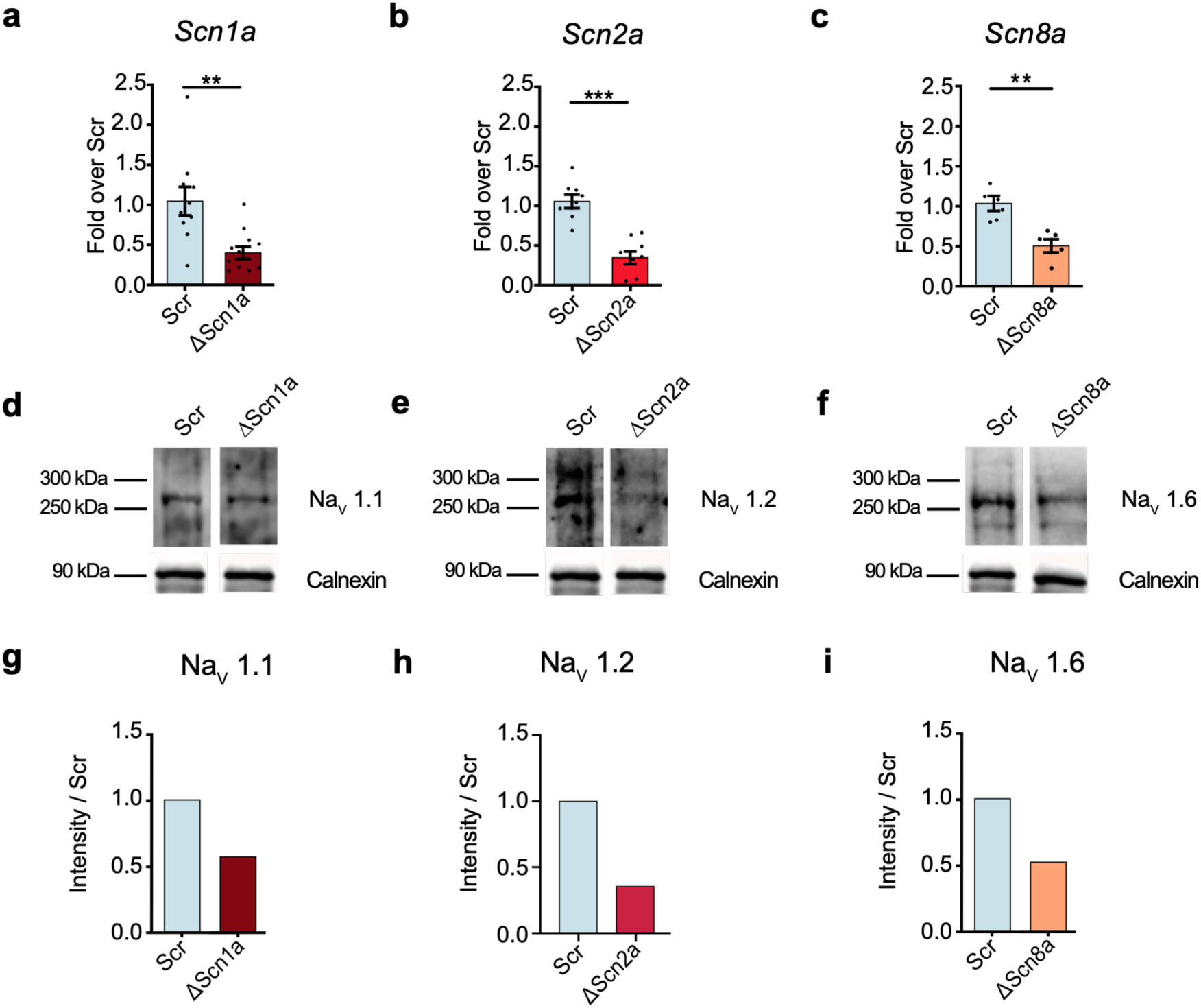
shRNA-mediated downregulation selectively reduces NaV subtypes expression. RT-qPCR analyses of **a.** *Scn1a* expression in ΔScn1a, **b.** *Scn2a* in ΔScn2a, and **c.** *Scn8a* in ΔScn8a neurons. Images of the Western blots of **d.** Na_V_ 1.1 in ΔScn1a, **e.** Na_V_ 1.2 in ΔScn2a, **f.** Na_V_1.6 in ΔScn8a neurons, and their respective quantifications in **g.**, **h.**, and **i.**

### Subtype-specific Na_V_ downregulation differentially affects neuronal survival and pro-inflammatory cytokine gene expression under basal conditions

We first investigated whether selective Na_V_ downregulation affected neuronal survival under basal conditions. Neurons were exposed to conditioned medium from unstimulated microglia (CM-Ctl), and the proportion of Map2^+^ cells was quantified at DIV10. ΔScn1a and ΔScn8a neurons displayed a modest but significant increase in the proportion of Map2^+^ cells in comparison to Scr controls (respectively 40.25%, p = 0.0215 and 41.39%, p = 0.0135 *vs.* 33.33% in Scr). In contrast, ΔScn2a neurons showed a reduced proportion of Map2^+^ cells (24.94%, p = 0.0126; **Fig. 3a, b**), indicating that reduction of Na_V_1.2 expression had a distinct effect on neuronal survival under basal conditions.

**Figure 3:**
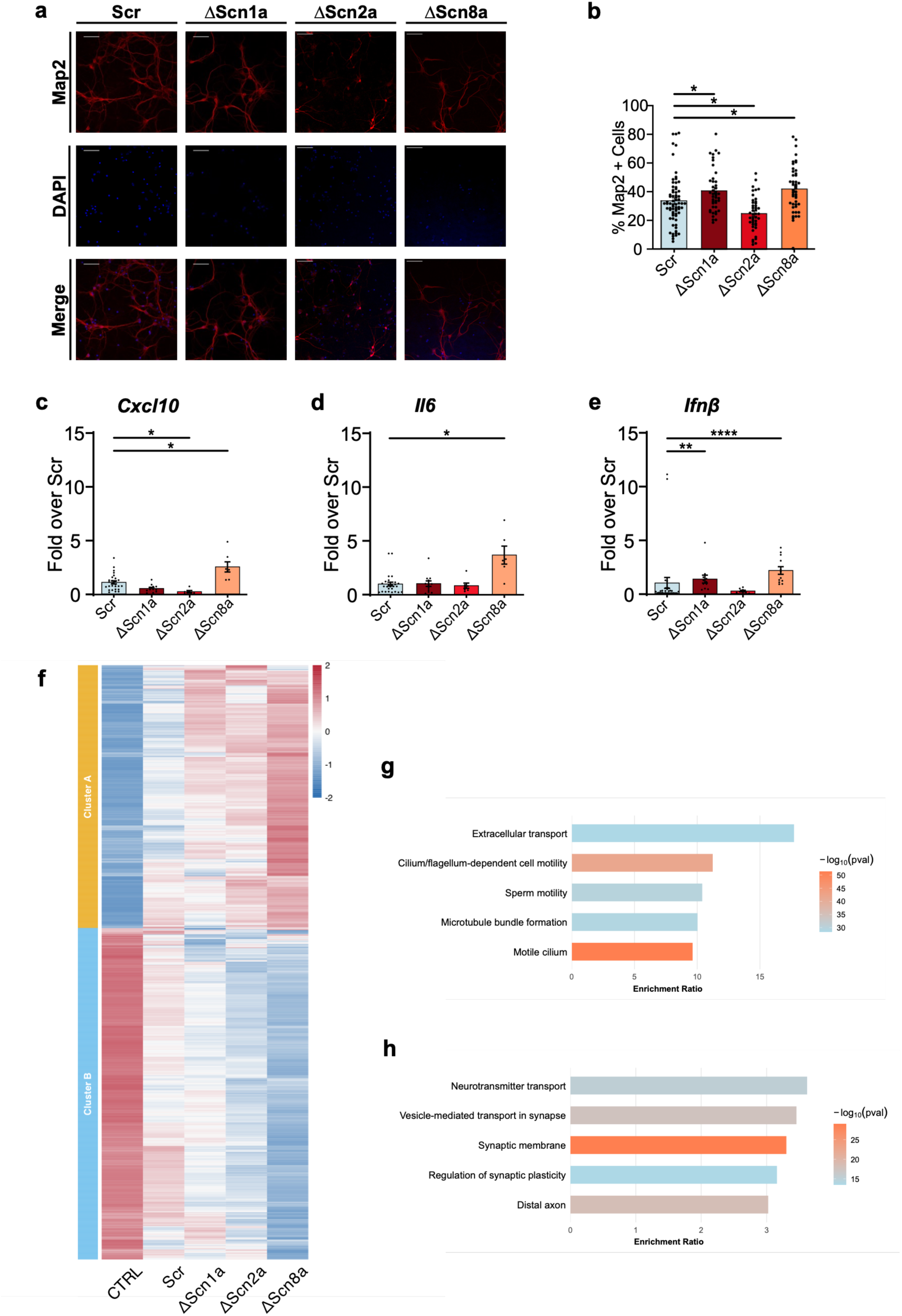
Na_V_ downregulations differentially affect neuronal survival and pro-inflammatory cytokine gene expression but do not reveal important transcriptomic disparities under basal conditions. **a.** Representative immunocytochemistry images for each condition: DAPI (blue), Map2 (red) **b.** Proportion of Map2^+^ cells over total DAPI^+^ nuclei normalized to Scr. RT-qPCR quantification of c. *Cxcl10*, d. *Il-6*, and e. *Ifn-ß* normalized to Scr **f.** Heatmap of the counts of the DEG identified between all conditions and CTRL that do not appear between Scr and CTRL (adj. pval < 0.05; FC > 2). Pathway enrichment analyses of genes included in **g.** cluster A, and **h.** cluster B. Figures display the top 5 enriched pathways ordered by enrichment ratio.

Because neurons can contribute to inflammatory signaling through the production of cytokines and other immune mediators (8–10), we next assessed how the selective downregulation of Na_V_ subtypes impacted the neuronal gene expression of inflammatory cytokine *Cxcl10* (**Fig. 3c**), *Il-6* (**Fig. 3d**) and *Ifn-β* (**Fig. 3e**). This analysis highlighted the distinctive consequences of each Na_V_ downregulation on the neurons’ inflammatory transcriptional profile. ΔScn8a neurons displayed 2.6-(p = 0.0341), 3.7-(p=0.0056), and 2.2-fold (p < 0.0001) increases in expression of all three cytokines (*Cxcl10*, *Il-6* and *Ifn-β* respectively) in comparison to Scr controls. In ΔScn1a neurons, only *Ifn-β* was increased by 1.38-fold (p = 0.0025), whereas ΔScn2a neurons showed a 0.23-fold (p = 0.0047) reduction in *Cxcl10* expression, with *Il-6* and *Ifn-β* levels remaining comparable to Scr. Overall, these results highlight that subtype-specific Na_V_ downregulations differentially shape the basal neuronal phenotype, with distinct effects on both neuronal survival and inflammatory gene expression.

### Subtype-specific Na_V_ downregulations affect the transcription of genes involved in neuronal and synaptic transport pathways

To further characterize the molecular consequences of Na_V_ downregulation, we performed bulk RNA sequencing (RNAseq) in ΔScn1a, ΔScn2a, and ΔScn8a neurons under basal conditions. Comparison of each Na_V_ knockdown with untransfected control neurons (Ctl) identified 423 differentially expressed genes (DEGs) in ΔScn1a (130 up, 293 downregulated), 666 in ΔScn2a (360 up, 306 downregulated), and 1818 in ΔScn8a (1006 up, 812 downregulated), resulting in a total of 1912 unique DEGs. Genes differentially expressed in the Scr *vs.* Ctl comparison (31 DEGs, 11 up, 20 downregulated), were excluded from subsequent analysis to account for the transcriptional effects associated with the viral construct.

Hierarchical clustering on the resulting 1881 genes, identified two major clusters based on their number of reads (**Fig. 3f**). Cluster A, comprising 827 genes, consisted predominantly of genes upregulated relative to Ctl neurons. Conversely, cluster B contained 1042 genes mostly downregulated in comparison to Ctl neurons. Pathway enrichment analysis revealed the enrichment in cluster A of pathways related to cilia and cellular transport (*e.g.* “Extracellular transport”, “Flagellum-dependent cell motility”, “Motile cilium” …; **Fig. 3g**), and in cluster B, of pathways related to synaptic transport and modulation (*e.g.* “Neurotransmitter transport”, “Synaptic membrane”, “Regulation of synaptic plasticity” …; **Fig. 3h**).

Overall, the differential downregulation of Na_V_s, in absence of any inflammatory stimulus, led to similar transcriptomic consequences, notably underlining an important modulation of pathways related to the synaptic machinery in neurons. However, these modifications were not correlated with changes in neuronal survival or inflammatory cytokine production.

### Inflammatory stimulus differentially affects neuronal survival following subtype-specific Na_V_ downregulation

We next investigated whether subtype-specific Na_V_ downregulation altered neuronal responses to an inflammatory stimulus. To induce an inflammatory environment, neurons were exposed for 24 h (from DIV9 to 10) to conditioned medium collected from LPS-activated microglia (CM-Inf). The addition of CM-inf led to a statistically significant decrease in the proportion of Map2^+^ cells in ΔScn1a + CM-inf neurons compared to Scr + CM-inf (28.48% *vs.* 38.78%, p = 0.0171). However, no statistical difference was observed in ΔScn2a + CM-inf (31.02%), and in ΔScn8a + CM-inf (36.78 %) compared with Scr + CM-inf cultures (**Fig. 4a, b**). This proportion did not differ either between Scr + CM-inf cultures and Ctl + CM-inf cultures (38.78% vs. 35.41%). Comparison with corresponding basal condition results further highlighted the specific effects of inflammatory stimulation on neurons with subtype-specific Na_V_ downregulations. These results notably underline the deleterious effect of inflammatory stimuli in ΔScn1a neurons.

**Figure 4:**
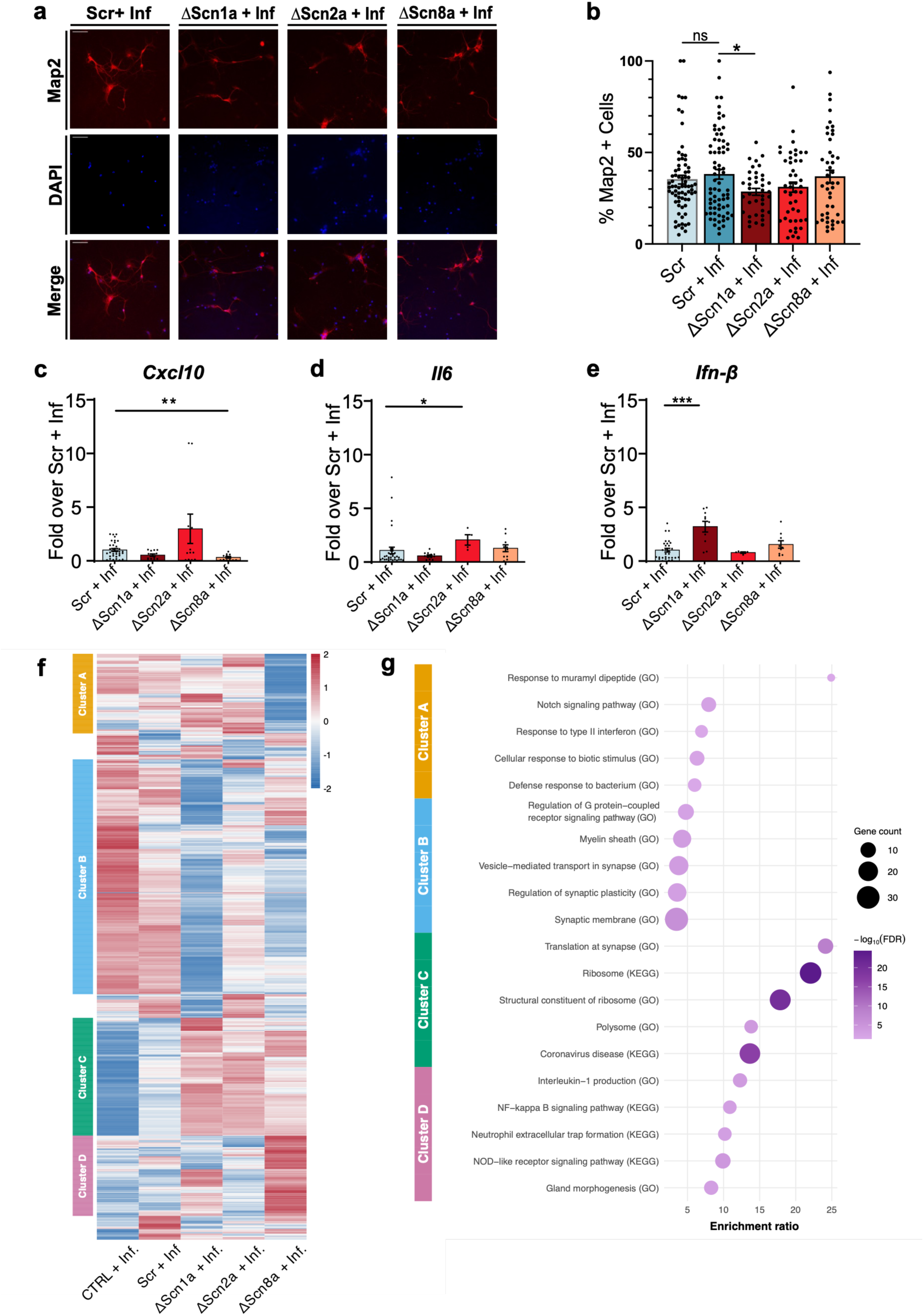
Inflammatory challenge unmasks divergent transcriptomic responses across subtype-specific downregulation of Na_V_ in neurons. **a.** Representative immunocytochemistry images for each condition: DAPI (blue), Map2 (red) **b.** Proportion of Map2^+^ cells over total DAPI^+^ nuclei normalized to Scr + inf. RT-qPCR quantification of c. *Cxcl10*, d. *Il-6*, and e. *Ifn-ß* normalized to Scr + Inf. **f.** Heatmap of the counts of the DEG identified between all conditions and CTRL + Inf. that do not appear between Scr + Inf. and CTRL + Inf. (adj. pval < 0.05; FC > 2). **g.** Pathway enrichment analyses of genes included in cluster A, B, C and D. The figure displays the top 5 enriched pathways ordered by enrichment ratio.

These results were however not correlated with an overall increase in *Cxcl10*, *Il-6* and *Ifn-β* expression but rather with distinct inflammatory profiles in each condition. ΔScn1a + CM-inf displayed a sole significant 3.11-fold increase in *Ifn-β* levels (p = 0.0006), ΔScn2a + CM-inf ’s response was limited to a 1.95-fold increase in *Il-6* (p = 0.0381), and ΔScn8a + CM-inf even showed a 0.27-fold decrease in *Cxcl10* (p = 0.0012) when compared to Scr + CM-inf.

### Subtype-specific Na_V_ downregulation led to different transcriptional responses to inflammation

To further investigate the molecular mechanisms underlying the differential neuronal responses to inflammatory stimulation, we performed bulk RNA sequencing following exposure to CM-Inf. Comparison of each condition after CM-Inf exposure to its corresponding basal condition identified 605 DEGs in ΔScn1a + CM-Inf *vs.* ΔScn1a, 92 DEGs in ΔScn2a + CM-Inf *vs.* ΔScn2a, and 106 DEGs in ΔScn8a + CM-Inf *vs.* ΔScn8a, and 1802 DEGs in Ctl + CM-Inf *vs.* Ctl, corresponding to 951 unique DEGs. As in the basal analysis, genes differentially expressed between Scr + CM-Inf and Ctl + CM-Inf (185 DEGs) were excluded to account for transcriptional effects associated with the viral construct.

Hierarchical clustering of the resulting 766 genes identified four major clusters labelled A to D (**Fig. 4f**), revealing substantially more divergent transcriptional profiles than those observed under basal conditions. Cluster A contained 104 genes that appeared strongly downregulated only in ΔScn8a + CM-inf, whereas cluster B comprised 307 genes predominantly downregulated in ΔScn1a + CM-inf, with other conditions showing more scattered expression profiles. Cluster C, comprising 154 genes, showed increased gene expression across all Na_V_ subtypes downregulations, while cluster D, containing 105 genes, was characterized by an important upregulation in ΔScn8a + CM-inf.

Pathway enrichment analysis further highlighted condition specific tendencies. Cluster A and D, strongly influenced by the ΔScn8a + CM-inf condition, were both enriched in inflammatory signaling pathways. Downregulated genes in cluster A were associated with pathways including “Response to type II interferon”, “Cellular response to biotic stimulus”, or “Defense response to bacterium”. Conversely, cluster D, upregulated in ΔScn8a + CM-inf, was associated with the enrichment of “NF-κB signaling pathway”, “Interleukin-1 production”, or “Neutrophil extracellular trap formation”. Cluster B, characterized by downregulated genes in ΔScn1a + CM-inf, was associated with pathways related to synaptic function (*e.g.* “Synaptic membrane”, “Regulation of synaptic plasticity”, “Vesicle-mediated transport in synapse” …). Finally, cluster C, commonly upregulated across all conditions Na_V_ subtype downregulations, was enriched for pathways related to protein synthesis, including “Ribosome”, “Structural constituent of ribosome”, and “Translation at synapse” (**Fig. 4g**).

Together, these results underline that neurons differentially respond to inflammation depending on the expression of Na_V_ subtypes. In particular, ΔScn8a neurons displayed prominent alterations in inflammatory signaling pathways, whereas ΔScn1a neurons showed greater modulation of synaptic pathways. These findings support the hypothesis that inflammatory stimuli, such as seizures, may have differential neuronal etiology-dependent consequences.

## Discussion

The high incidence of epilepsy and important proportion of drug-resistant patients make it a major health concern despite the large number of anti-seizure medications available. Although the implication of Na_V_ subtypes in epileptic syndromes and the relationship between seizures and inflammation have been extensively described, the differential implications of inflammation in neurons presenting with Na_V_ LoF mutations remain largely unexplored. In the present study, we investigated whether the downregulation of either *Scn1a*, *Scn2a*, *or Scn8a* led to different survival, inflammatory or transcriptional outcomes in the presence or absence of an inflammatory stimulus, in mouse neuronal cortical cultures. The primary outcome of our investigation was that subtype-specific Na_V_ downregulation did not produce a uniform phenotype. Rather, differences that appear minor under basal conditions led to important discrepancies following exposure to an inflammatory stimulus.

Under basal conditions, Na_V_ downregulation produced relatively modest but subtype-dependent effects on neuronal survival and inflammatory gene expression. In particular, in contrast to ΔScn1a and ΔScn8a conditions, ΔScn2a neurons displayed a distinct basal neuronal vulnerability rather than a distinct inflammatory or synaptic profile. The reduction in MAP2+ cells following *Scn2a* downregulation is consistent with a role for Na_V_1.2 in neuronal survival during development. Similar results have previously been reported in mouse *Scn2a*^-/-^ models with a noticeable impact on neuronal apoptosis in the brainstem, contributing to the premature death of the animals (29). However, more recent studies in mouse and human organoids on *Scn2a* haploinsufficiency highlight the over-pruning of synapses by microglia during development rather than neuronal loss (30,31). The apparent augmentation in the proportion of neurons in ΔScn1a and ΔScn8a conditions does not seem to have been previously reported and must be taken with caution. The simultaneous increase in *Ifn-β* signaling in both conditions could point to the protective effects of *Ifn-β* on neural stem-cells, leading to larger neuronal populations at DIV 10 (32). These findings are consistent with the idea that the contribution of individual Na_V_ subtypes to neuronal homeostasis is not interchangeable. Reduced neuronal survival after *Scn2a* downregulation could reflect altered excitability, impaired maturation, or secondary effects on neuronal homeostasis.

The most informative results of our study emerged after the addition of an inflammatory stimulus, in the form of LPS-stimulated microglia culture medium, to ΔScn1a and ΔScn8a neurons. In contrast to the relatively limited effects observed under basal conditions, inflammatory stimulation revealed more pronounced and qualitatively distinct responses across conditions. This observation is particularly relevant in the context of epilepsy, where inflammatory signaling and seizures form a reciprocal relationship influencing neuronal excitability, synaptic function, and neuronal damage (7,33,34).

Among the three subtypes, *Scn8a* downregulation produced the most prominent inflammatory phenotype. Under basal conditions, ΔScn8a neurons showed increased expression of *Cxcl10*, *Il-6*, and *Ifn-β*, while inflammatory stimulation was associated with marked remodeling of transcriptomic programs related to immune-related pathways. To that extent, the downregulation of Interferon and *TNF* signaling and simultaneous upregulation of *Il-1β* and *NF-kB* signaling highlight a profound remodeling of the inflammatory landscape in these neurons. These results echo previous work implicating Na_V_ 1.6 in inflammatory and neurodegenerative processes. In a mouse model of multiple sclerosis, Na_V_ 1.6 was reported to promote inflammation and axonal degeneration following demyelination, and suppressed expression of *Scn8a* attenuated disease-associated pathology (35). Na_V_ 1.6 function therefore appears closely linked to the remodeling of neuronal inflammatory signaling in response to neuroinflammation. Whether this remodeling reflects a broader dysregulation of inflammatory signaling, as in our data, or a predominantly pro-inflammatory response, as previously reported, may therefore be context dependent.

The response of ΔScn1a neurons to inflammation was functionally different and was associated with a specific inflammatory signature. Specifically, ΔScn1a neurons here displayed a profound downregulation of pathways related to synaptic membranes, neuron-to-neuron synapses, and postsynaptic specialization, highlighting the profound remodeling occurring at the synapse. Similar effects were already commonly apparent under basal conditions across Na_V_ downregulations. Inflammatory stress, therefore, seems to amplify downstream synaptic consequences of reduced Na_V_1.1. This result is of particular interest in the context of DS, in which reduced Na_V_1.1 function preferentially compromises inhibitory interneuron excitability and thereby disrupts network inhibition (36,37). Further, previous studies in DS mouse models have highlighted synaptic alterations, notably reductions in the number of inhibitory synapses, and alterations in their vesicular release probability (38,39). However, because the present experiments were performed in mixed cortical cultures and bulk RNAseq was used, the observed transcriptional changes cannot be assigned specifically to inhibitory interneurons. The data therefore support an association between *Scn1a* downregulation and altered synaptic programs under inflammatory stress, rather than a definitive interneuron-specific mechanism. This *Scn1a*-specific result could further explain the anticipated role of seizures and neuroinflammatory processes in the onset and debilitating progression of both epileptic and developmental aspects of DS. It was notably previously shown that the induction of short, repeated seizures, could lead to the acquisition of a severe DS-like phenotype in mice carrying a *Scn1a* variant responsible for an originally mild phenotype (40).

Several limitations should be considered when interpreting the results of our study. First, shRNA-mediated reduction of Na_V_ expression does not reproduce the molecular consequences of a specific pathogenic *SCN1A*, *SCN2A*, or *SCN8A* variant and may not capture variant-specific effects on channel functionality. Second, the primary cortical cultures contain multiple neuronal populations; therefore, bulk transcriptomic changes cannot be assigned to defined excitatory or inhibitory cell types. Most importantly, our cultures include a reduced but still present population of astrocytes. Because astrocytes can respond directly to inflammatory stimuli, their contribution to the bulk transcriptomic profiles cannot be excluded. However, astrocytes do not commonly express Na_V_ 1.1, Na_V_ 1.2 or Na_V_ 1.6. Therefore, their relative contribution is unlikely for the subtype-specific differences observed here. These limitations define rather than diminish the most informative next steps: determining whether the subtype-specific transcriptional responses are linked to distinct changes in neuronal excitability, validating candidate signaling mechanisms at the protein level, and testing whether the same relationships are preserved in genetically precise disease models.

## Conclusion

In conclusion, our findings indicate that Na_V_1.1, Na_V_1.2, and Na_V_1.6 downregulations are associated with distinct neuronal responses to inflammatory stress. Na_V_1.1 reduction was preferentially associated with synaptic transcriptional changes, targeting Na_V_1.2 resulted in a distinct basal vulnerability, and Na_V_1.6 downregulation displayed the most prominent inflammatory transcriptional signature. These observations suggest that the biological consequences of Na_V_ dysfunction are not determined solely by their canonical role in membrane excitability but may also depend on subtype-specific responses to cellular stress. They notably shed light on the relevance of inflammatory events in the onset and progression of epileptic syndromes related to Na_V_ loss-of-function variants.

## Author Contributions

D.J., D.G., N.H., S.M., A.M.K.: study concept and design. D.J., D.G., N.H., S.M.: data acquisition and analysis. D.J., D.G., S.M., A.M.K.: drafting manuscript

## Supporting information

Supplementary Figure 1: Cytokine Array of CM-Inf

## Acknowledgments

Graphical abstract and schematic representations were created using BioRender. We thank Yanyan Wang and Hariprasath Ragupathy for their technical assistance.

## Ethical approval

We confirm that we have read the Journal’s position on issues involved in ethical publication and affirm that this report is consistent with those guidelines. All animal experiments were carried out in accordance with the national ethic principles (registration no. T0344/12, State Office for Health (LAGeSo), Berlin, Germany).

## References

1. GBD Epilepsy Collaborators. Global, regional, and national burden of epilepsy, 1990-2021: a systematic analysis for the Global Burden of Disease Study 2021. Lancet Public Health. 2025 Mar;10(3):e203–27. doi:10.1016/S2468-2667(24)00302-5 PubMed PMID: 40015291; PubMed Central PMCID: PMC11876103.

2. Zhang MW, Liang XY, Wang J, Gao LD, Liao HJ, He YH, et al. Epilepsy-associated genes: an update. Seizure Eur J Epilepsy. 2024 Mar;116:4–13. doi:10.1016/j.seizure.2023.09.021

3. Brunklaus A, Lal D. Sodium channel epilepsies and neurodevelopmental disorders: from disease mechanisms to clinical application. Dev Med Child Neurol. 2020 Jul;62(7):784–92. doi:10.1111/dmcn.14519 PubMed PMID: 32227486.

4. Jenkins PM, Bender KJ. Axon initial segment structure and function in health and disease. Physiol Rev. 2025 Apr 1;105(2):765–801. doi:10.1152/physrev.00030.2024 PubMed PMID: 39480263; PubMed Central PMCID: PMC12239863.

5. Auvin S, Shin D, Mazarati A, Nakagawa J, Miyamoto J, Sankar R. Inflammation exacerbates seizure-induced injury in the immature brain. Epilepsia. 2007;48 Suppl 5:27–34. doi:10.1111/j.1528-1167.2007.01286.x PubMed PMID: 17910578.

6. Choi J, Nordli DR, Alden TD, DiPatri A, Laux L, Kelley K, et al. Cellular injury and neuroinflammation in children with chronic intractable epilepsy. J Neuroinflammation. 2009 Dec 19;6:38. doi:10.1186/1742-2094-6-38 PubMed PMID: 20021679; PubMed Central PMCID: PMC2811703.

7. Villasana-Salazar B, Vezzani A. Neuroinflammation microenvironment sharpens seizure circuit. Neurobiol Dis. 2023 Mar;178:106027. doi:10.1016/j.nbd.2023.106027 PubMed PMID: 36736598.

8. Knoblach SM, Fan L, Faden AI. Early neuronal expression of tumor necrosis factor-alpha after experimental brain injury contributes to neurological impairment. J Neuroimmunol. 1999 Mar 1;95(1–2):115–25. doi:10.1016/s0165-5728(98)00273-2 PubMed PMID: 10229121.

9. Suzuki S, Tanaka K, Nagata E, Ito D, Dembo T, Fukuuchi Y. Cerebral neurons express interleukin-6 after transient forebrain ischemia in gerbils. Neurosci Lett. 1999 Mar 5;262(2):117–20. doi:10.1016/s0304-3940(99)00051-8 PubMed PMID: 10203245.

10. Wood LB, Singer AC. Neurons as Immunomodulators: From Rapid Neural Activity to Prolonged Regulation of Cytokines and Microglia. Annu Rev Biomed Eng. 2025 May;27(1):55–72. doi:10.1146/annurev-bioeng-110122-120158 PubMed PMID: 39805040; PubMed Central PMCID: PMC12486157.

11. Zipp F, Bittner S, Schafer DP. Cytokines as emerging regulators of central nervous system synapses. Immunity. 2023 May 9;56(5):914–25. doi:10.1016/j.immuni.2023.04.011 PubMed PMID: 37163992; PubMed Central PMCID: PMC10233069.

12. De Simoni MG, Perego C, Ravizza T, Moneta D, Conti M, Marchesi F, et al. Inflammatory cytokines and related genes are induced in the rat hippocampus by limbic status epilepticus. Eur J Neurosci. 2000 Jul;12(7):2623–33. doi:10.1046/j.1460-9568.2000.00140.x PubMed PMID: 10947836.

13. Turrin NP, Rivest S. Innate immune reaction in response to seizures: implications for the neuropathology associated with epilepsy. Neurobiol Dis. 2004 Jul;16(2):321–34. doi:10.1016/j.nbd.2004.03.010 PubMed PMID: 15193289.

14. Victor TR, Hage Z, Tsirka SE. Prophylactic Administration of Cannabidiol Reduces Microglial Inflammatory Response to Kainate-Induced Seizures and Neurogenesis. Neuroscience. 2022 Sep 15;500:1–11. doi:10.1016/j.neuroscience.2022.06.010 PubMed PMID: 35700815.

15. Alonso C, Satta V, Hernández-Fisac I, Fernández-Ruiz J, Sagredo O. Disease-modifying effects of cannabidiol, β-caryophyllene and their combination in Syn1-Cre/Scn1aWT/A1783V mice, a preclinical model of Dravet syndrome. Neuropharmacology. 2023 Oct 1;237:109602. doi:10.1016/j.neuropharm.2023.109602

16. Cha J, Filatov G, Smith SJ, Gammaitoni AR, Lothe A, Reeder T. Fenfluramine increases survival and reduces markers of neurodegeneration in a mouse model of Dravet syndrome. Epilepsia Open. 2024 Feb;9(1):300–13. doi:10.1002/epi4.12873 PubMed PMID: 38018342; PubMed Central PMCID: PMC10839300.

17. Yang Y, Wang B, Zhong Z, Chen H, Ding W, Hoi MPM. Clonazepam attenuates neurobehavioral abnormalities in offspring exposed to maternal immune activation by enhancing GABAergic neurotransmission. Biochem Pharmacol. 2021 Oct;192:114711. doi:10.1016/j.bcp.2021.114711 PubMed PMID: 34324871.

18. Qiao X, Werkman TR, Gorter JA, Wadman WJ, van Vliet EA. Expression of sodium channel α subunits 1.1, 1.2 and 1.6 in rat hippocampus after kainic acid-induced epilepsy. Epilepsy Res. 2013 Sep;106(1–2):17–28. doi:10.1016/j.eplepsyres.2013.06.006 PubMed PMID: 23886654.

19. Beaudoin GMJ, Lee SH, Singh D, Yuan Y, Ng YG, Reichardt LF, et al. Culturing pyramidal neurons from the early postnatal mouse hippocampus and cortex. Nat Protoc. 2012 Sep;7(9):1741–54. doi:10.1038/nprot.2012.099 PubMed PMID: 22936216.

20. Platow RJ, Pommer S, Brauer J, Wang Y, Mani S, Kaindl AM. Microglial activation is inhibited by selective anti-seizure medications. Inflamm Res. 2025 Aug 23;74(1):112. doi:10.1007/s00011-025-02076-7

21. Schindelin J, Arganda-Carreras I, Frise E, Kaynig V, Longair M, Pietzsch T, et al. Fiji: an open-source platform for biological-image analysis. Nat Methods. 2012 Jul;9(7):676–82. doi:10.1038/nmeth.2019

22. Elizarraras JM, Liao Y, Shi Z, Zhu Q, Pico AR, Zhang B. WebGestalt 2024: faster gene set analysis and new support for metabolomics and multi-omics. Nucleic Acids Res. 2024 Jul 5;52(W1):W415–21. doi:10.1093/nar/gkae456

23. R Core Team. R: A language and environment for statistical computing [Internet]. R Foundation for Statistical Computing; 2024 [cited 2026 Aug 23]. Available from: https://zenodo.org/doi/10.5281/zenodo.21842137 doi:10.5281/ZENODO.21842137

24. Wickham H. ggplot2: elegant graphics for data analysis. Second edition. Cham: Springer international publishing; 2016. 1 p. (Use R!).

25. Kolde R. pheatmap: Pretty Heatmaps [Internet]. 2010 [cited 2026 Aug 23]. p. 1.0.13. Available from: https://CRAN.R-project.org/package=pheatmap doi:10.32614/CRAN.package.pheatmap

26. Moutin E, Hemonnot AL, Seube V, Linck N, Rassendren F, Perroy J, et al. Procedures for Culturing and Genetically Manipulating Murine Hippocampal Postnatal Neurons. Front Synaptic Neurosci. 2020 Apr 30;12. doi:10.3389/fnsyn.2020.00019

27. Dutton SB, Makinson CD, Papale LA, Shankar A, Balakrishnan B, Nakazawa K, et al. Preferential inactivation of Scn1a in parvalbumin interneurons increases seizure susceptibility. Neurobiol Dis. 2013 Jan;49:211–20. doi:10.1016/j.nbd.2012.08.012 PubMed PMID: 22926190; PubMed Central PMCID: PMC3740063.

28. Yamagata T, Ogiwara I, Tatsukawa T, Suzuki T, Otsuka Y, Imaeda N, et al. Scn1a-GFP transgenic mouse revealed Nav1.1 expression in neocortical pyramidal tract projection neurons. eLife. 2023 May 23;12:e87495. doi:10.7554/eLife.87495 PubMed PMID: 37219072; PubMed Central PMCID: PMC10205085.

29. Planells-Cases R, Caprini M, Zhang J, Rockenstein EM, Rivera RR, Murre C, et al. Neuronal Death and Perinatal Lethality in Voltage-Gated Sodium Channel αII-Deficient Mice. Biophys J. 2000 Jun;78(6):2878–91. doi:10.1016/S0006-3495(00)76829-9

30. Wu J, Chen X, Zhang J, Wettschurack K, Robinson M, Li W, et al. Human microglia in brain assembloids display region-specific diversity and respond to hyperexcitable neurons carrying *SCN2A* mutation. Sci Adv. 2026 Feb 20;12(8):eady2977. doi:10.1126/sciadv.ady2977

31. Wu J, Zhang J, Chen X, Wettschurack K, Que Z, Deming BA, et al. Microglial over-pruning of synapses during development in autism-associated SCN2A-deficient mice and human cerebral organoids. Mol Psychiatry. 2024 Aug;29(8):2424–37. doi:10.1038/s41380-024-02518-4

32. Hirsch M, Knight J, Tobita M, Soltys J, Panitch H, Mao-Draayer Y. The effect of interferon-β on mouse neural progenitor cell survival and differentiation. Biochem Biophys Res Commun. 2009 Oct;388(2):181–6. doi:10.1016/j.bbrc.2009.07.073

33. Conroy SM, Nguyen V, Quina LA, Blakely-Gonzales P, Ur C, Netzeband JG, et al. Interleukin-6 produces neuronal loss in developing cerebellar granule neuron cultures. J Neuroimmunol. 2004 Oct;155(1–2):43–54. doi:10.1016/j.jneuroim.2004.06.014

34. Gross A, Benninger F, Madar R, Illouz T, Griffioen K, Steiner I, et al. Toll-like receptor 3 deficiency decreases epileptogenesis in a pilocarpine model of SE -induced epilepsy in mice. Epilepsia. 2017 Apr;58(4):586–96. doi:10.1111/epi.13688

35. Alrashdi B, Dawod B, Schampel A, Tacke S, Kuerten S, Marshall JS, et al. Nav1.6 promotes inflammation and neuronal degeneration in a mouse model of multiple sclerosis. J Neuroinflammation. 2019 Nov 13;16(1):215. doi:10.1186/s12974-019-1622-1 PubMed PMID: 31722722; PubMed Central PMCID: PMC6852902.

36. Ogiwara I, Miyamoto H, Morita N, Atapour N, Mazaki E, Inoue I, et al. Nav1.1 Localizes to Axons of Parvalbumin-Positive Inhibitory Interneurons: A Circuit Basis for Epileptic Seizures in Mice Carrying an Scn1a Gene Mutation. J Neurosci. 2007 May 30;27(22):5903–14. doi:10.1523/JNEUROSCI.5270-06.2007 PubMed PMID: 17537961; PubMed Central PMCID: PMC6672241.

37. Liautard C, Scalmani P, Carriero G, De Curtis M, Franceschetti S, Mantegazza M. Hippocampal hyperexcitability and specific epileptiform activity in a mouse model of D ravet syndrome. Epilepsia. 2013 Jul;54(7):1251–61. doi:10.1111/epi.12213

38. Uchino K, Kawano H, Tanaka Y, Adaniya Y, Asahara A, Deshimaru M, et al. Inhibitory synaptic transmission is impaired at higher extracellular Ca2+ concentrations in Scn1a+/− mouse model of Dravet syndrome. Sci Rep. 2021 May 20;11(1):10634. doi:10.1038/s41598-021-90224-4

39. Almog Y, Mavashov A, Brusel M, Rubinstein M. Functional Investigation of a Neuronal Microcircuit in the CA1 Area of the Hippocampus Reveals Synaptic Dysfunction in Dravet Syndrome Mice. Front Mol Neurosci. 2022;15:823640. doi:10.3389/fnmol.2022.823640 PubMed PMID: 35370551; PubMed Central PMCID: PMC8966673.

40. Salgueiro-Pereira AR, Duprat F, Pousinha PA, Loucif A, Douchamps V, Regondi C, et al. A two-hit story: Seizures and genetic mutation interaction sets phenotype severity in SCN1A epilepsies. Neurobiol Dis. 2019 May;125:31–44. doi:10.1016/j.nbd.2019.01.006 PubMed PMID: 30659983.

