## Supplementary Figure 1: Cytokine Array of CM-Inf for "Subtype-specific downregulation of voltage-gated sodium channels shapes neuronal responses to neuroinflammation"

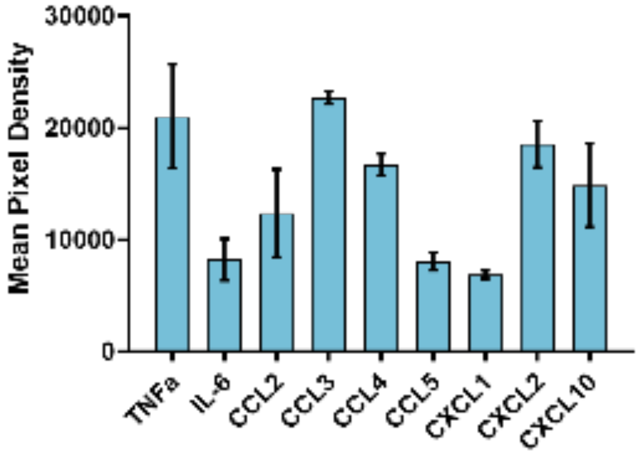


**Supplementary Figure 1: Cytokine Array of CM-Inf**
Microglia culture medium after 24h of LPS 10µM stimulation; n=3
